# Differences in the rare variant spectrum among human populations

**DOI:** 10.1101/063578

**Authors:** Iain Mathieson, David Reich

## Abstract

Mutations occur at vastly different rates across the genome, and populations, leading to differences in the spectrum of segregating polymorphisms. Here, we investigate variation in the rare variant spectrum in a sample of human genomes representing all major world populations. We find at least two distinct signatures of variation. One, consistent with a previously reported signature is characterized by an increased rate of TCC>TTC mutations in people from Western Eurasia and South Asia, likely related to differences in the rate, or efficiency of repair, of damage due to deamination of methylated guanine. We describe the geographic extent of this signature and show that it is detectable in the genomes of ancient, but not archaic humans. The second signature is private to certain Native American populations, and is concentrated at CpG sites. We show that this signature is not driven by differences in the CpG mutation rate, but is a result of the fact that highly mutable CpG sites are more likely to undergo multiple independent mutations across human populations, and the spectrum of such mutations is highly sensitive to recent demography. Both of these effects dramatically affect the spectrum of rare variants across human populations, and should be taken into account when using mutational clocks to make inference about demography.

## Introduction

For a process that provides such a fundamental contribution to genetic diversity, the human germline mutation rate is surprisingly poorly understood. Different estimates of the absolute mutation rate–the mean number of mutations per-generation, or per-year–are largely inconsistent with each other [1, 2], and similar uncertainty surrounds parameters such as the paternal age effect [3–5], the effect of life-history traits [6, 7], and the sequence-context determinants of mutations [5, 8]. Here, we investigate a related question. Rather than trying to determine the absolute values of parameters of the mutation rate, we ask how much the mutation spectrum–specifically, the relative rate of different classes of mutations–varies between different human populations. Because we are limited in our ability to observe the mutation rate directly (for example through studies of *de novo* mutations), we use the spectrum of segregating variation as a proxy. However, the relationship between mutation spectrum and variation spectrum can be affected by many factors, including selection, demography, recombination and gene conversion.

At least one class of polymorphism, most clearly represented by the trinucleotide mutation TCC>TTC but apparently including other classes as well, is known to be enriched in Europeans relative to East Asians and Africans [8, 9]. However the geographical extent, history, and biological basis for this signal are unclear. Analysis of tumor genomes has demonstrated a number of different mutational signatures operating at different rates in somatic cells and cancers, many of which can be linked to specific biological processes or environmental exposures [10–12]. It seems plausible that population-specific genetic factors of environmental exposures might similarly lead to variation in germline mutation spectra. Therefore, we used a dataset of high coverage genomes, representing much of the genetic diversity in present-day humans, to investigate the following three questions. First, is there evidence of any other differences in the spectrum of segregating variation across the world? Second, are these differences in variation driven by differences in mutation rates? Finally, if so, can we infer anything about the biological processes driving these differences?

## Results

We first analyzed data from 300 individuals sequenced to high coverage (mean coverage depth 43X) as part of the Simons Genome Diversity project [13] (SGDP). We classified single nucleotide polymorphisms (SNPs) into one of 96 mutational classes according to the SNP, and the two flanking bases. We represent these by the ancestral sequence and the derived base so for example “ACG>T” represents the ancestral sequence 3’–ACG– 5’ mutating to 3’–ATG–5’. We first focused on variants where there were exactly two copies of the derived allele in the entire sample of 300 individuals (we call these *f*_2_ variants or doubletons). This increases power to detect population-specific variation because rare variants tend to be recent mutations and are therefore highly differentiated between populations [14]. For each individual, we counted the number of *f*_2_ mutations in each mutational class that they carried, and normalized by the number of ATA>C mutations (the most common class and one that did not seem to vary across populations in a preliminary analysis). The normalized mutation intensities form a 96×300 matrix, and we used non-negative matrix factorization [11, 15] (NMF, implemented in the *NMF* package [16] in R) to identify specific mutational features. NMF decomposes a matrix into a set of sparse factors, here putatively representing different mutational processes, and individual-specific loadings for each factor, measuring the intensity of each process in each individual. It has been used extensively in the analysis of somatic mutations in cancer genomes [10, 11, 17, 18]. An advantage over PCA is that NMF tends to provide components that are sparser and more interpretable.

NMF requires us to specify the number of signatures (the factorization rank) in advance. For *f*_2_ variants we chose a factorization rank of 4, based on standard diagnostic criteria (Supplementary Fig. 1). This identified four mutational signatures; of which two were uncorrelated with each other, were robust across frequencies, replicated in non-cell-line samples, were consistent across samples from the same populations, and had clear geographic distributions (Figure 1, Supplementary Fig. 2). Signature 1 corresponds to the previously described European signal [8] characterized by TCC>T, ACC>T, CCC>T and TCT>T (possibly also including CCG>T, which overlaps with signature 2). Loadings of this component almost perfectly separate West Eurasians from other populations, with South-West Asians intermediate. It is seen most strongly in Western and Mediterranean Europe, with decreasing intensity in Northern and Eastern Europe, the Near East and South-west Asia. The COSMIC catalog of somatic mutation in cancer [19] is a database of mutational signatures extracted from samples of tumor genomes, also using NMF. Comparing with all the COSMIC signatures, we found that our signature 1 is most similar to COSMIC signature 11 (Pearson correlation *ρ*=0.81) which is most commonly found in melanoma and glioblastoma and is associated with use of chemotherapy drugs which act as alkylating agents, damaging DNA through guanine methylation.

**Figure 1:**
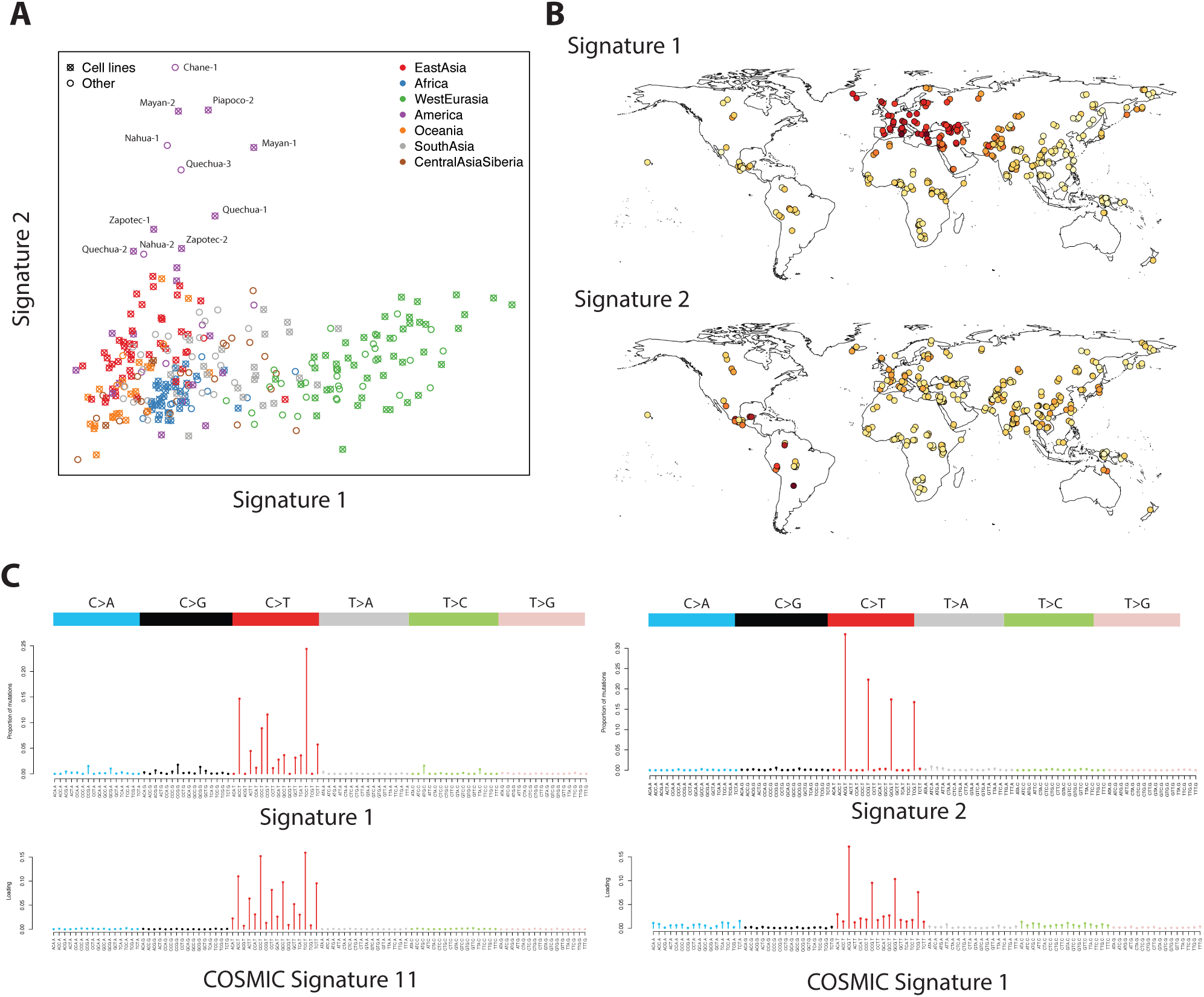
Distribution and characterization of signatures 1 and 2 for *f*_2_variants. **A**: Factor coefficients for these two signatures, for 300 individual samples colored by region. **B**: Geographic representation of the factor loadings from panel **A**. Darker colors represent higher loadings. **C**: Characterization of the signatures in terms of mutation intensity for each of 96 possible classes. Bars are scaled by the frequency of each trinucleotide in the human reference genome. Below, the most highly correlated signatures from the COSMIC database are shown for comparison.

Signature 2 is restricted to some South and Central American populations and, possibly, Aboriginal Australians. It is characterized by NCG>T mutations similar to the signature caused by deamination of methylated cytosine at CpG sites, corresponding to COSMIC signature 1 (*ρ*=0.96). Interestingly, this signal is found in South America in Andean populations like Quechua and Piapoco, and in Central American populations such as Mayan and Nahua, but not in the closely related Amazonian Surui and Karitiana, nor in North American populations.

The remaining two signatures are more difficult to characterize (Supplementary Fig. 2). Signature 3 is characterized by GT>GG mutations, particularly GTG>GGG. It is found in some East Asian and some South American populations but is not consistent within populations. For example, it is strongest in one Han sample (B_Han-3), but not at all increased in the two other samples from the same population. All affected samples are derived from cell lines. It does not match any mutational signature seen in COSMIC (maximum *ρ*=0.16). Plausibly this represents some as-yet uncharacterized cell-line artifact, or a very localized difference in mutation process. Signature 4 affects almost all mutation types, possibly representing a background mutation spectrum, and is most correlated with COSMIC signature 5 (*ρ*=0.60) which is found in all cancers and has unknown aetiology. It is significantly reduced in only a single cell-line derived sample (S_Quechua-2), so probably represents some unidentified cell-line or data processing artifact.

We checked whether these effects could be detected in singletons. At *f*_1_the variation is apparently dominated by cell line artifacts because principal component analysis (PCA) separates cell line from non cell line derived samples (Supplementary Fig. 3A). However, NMF on *f*_1_variants excluding cell line derived samples recovers signatures consistent with signatures 1 and 2 (Supplementary Fig. 3B-C), although it does not substantially separate out Native Americans based on signature 2. PCA on *f*_2_ variants does not distinguish cell line samples, but does separate samples by geographic region, and recovers factor loadings consistent with NMF-derived signatures 1-3 (Supplementary Fig. 4). To check that our results were not an artifact of the normalization we used, we repeated the analysis normalizing by the total number of mutations in each sample, rather than the number of ATA>C mutations, and obtained equivalent results (Supplementary Figure 5).

We replicated these results using data from phase 3 of the 1000 Genomes project [20] (Methods). To do this, we counted *f*_2_ and *f*_3_variants in each trinucleotide class and then, for each individual, computed the proportion of the total mutations carried by that individual that were in each of signatures 1 and 2 (Figure 2). This confirmed that that mutations consistent with signature 1 are enriched in populations of European and South Asian ancestry (Figure 2A; mean proportions 0.085, 0.077; Z-score for difference *Z*=42; for European/South Asian compared to all other populations) and that mutations consistent with signature 2 are enriched in Peruvians in Lima (PEL) and people of Mexican ancestry in Los Angeles (MXL) – the two 1000 Genomes populations with the most Native American ancestry (Figure 2B; mean proportions 0.216, 0.172; *Z*=34 for PEL+MXL compared to all other populations). Thus, the observed differences in the spectrum of variation are consistent across datasets. We then asked whether these differences could be interpreted as differences in the mutational spectrum.

**Figure 2:**
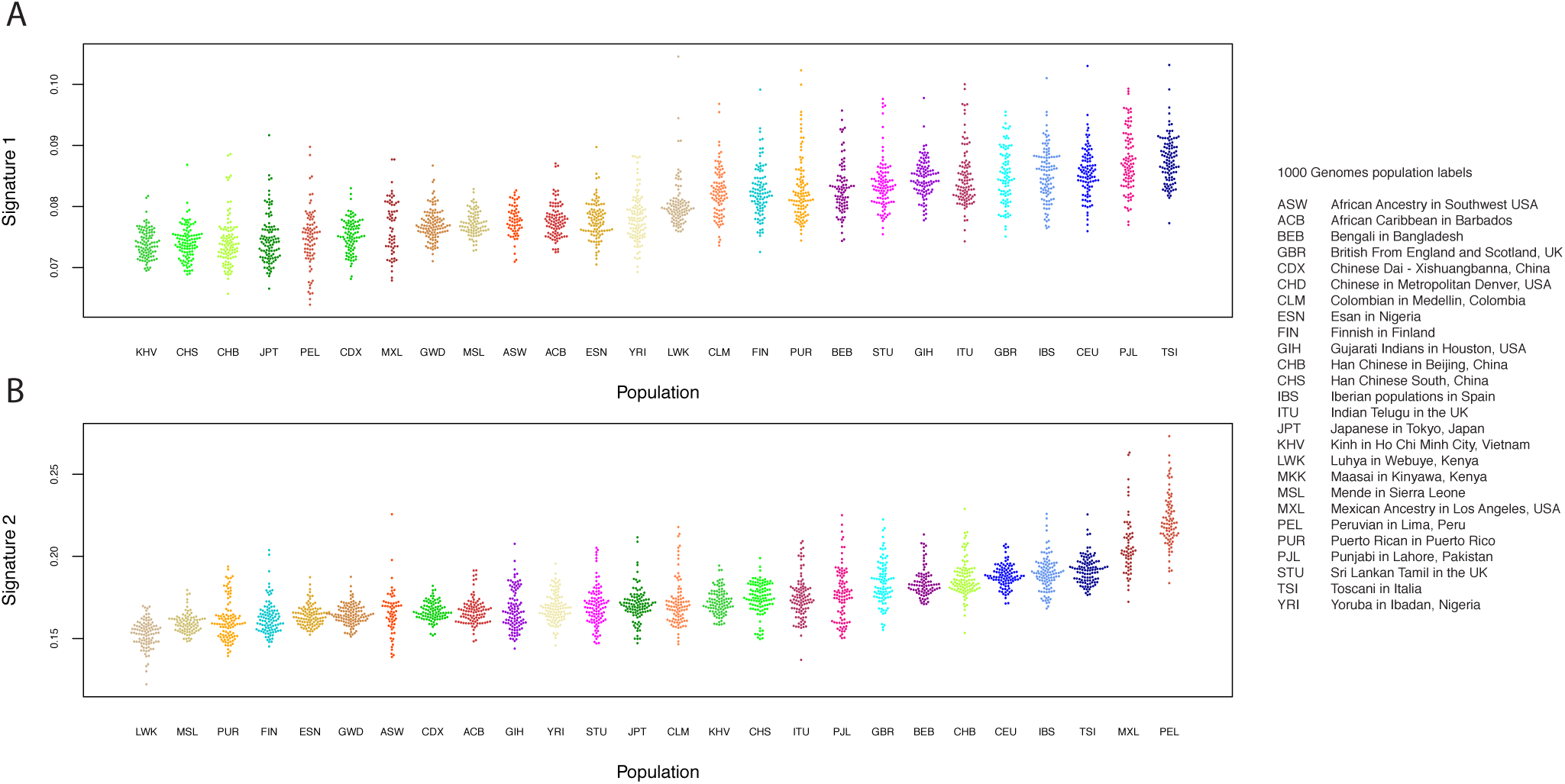
Signatures 1 and 2 in the 1000 Genomes. **A**: Proportions of *f*_2_ and *f*_3_ variants in signature 1 (here defined as TCT>T, TCC>T, CCC>T and ACC>T) in each 1000 Genomes individual, by population. **B**: Proportions of *f*_2_ and *f*_3_ variants in signature 2 (here defined as NCG>T, for any N) in each 1000 Genomes individual, by population (five outlying samples excluded).

To investigate whether non-mutational processes could be driving these differences, we first investigated the dependence of the two signatures on four genomic features. First we investigated dependence on transcriptional strand by classifying each mutation (not collapsed with its reverse complement, and defined on the + strand) according to whether it was on the coding or noncoding strand obtained from the UCSC genome browser (Methods). Signature 1 shows a skew whereby the C>T mutation is more likely to occur on the transcribed (i.e. noncoding) strand in West Eurasians, relative to populations from other regions (Figure 3 A&B). Because transcription coupled repair is more likely to repair mutations on the transcribed strand [21] this result, consistent with Harris (2015) [8], suggests that the excess signature 1 mutations in West Eurasians are driven more by G>A than by C>T mutations. Signature 2 shows a global skew where the C>T mutation is more likely to occur on the untranscribed strand, consistent with these mutations resulting from deamination of methylated cytosine, and we do not see a significant difference between individuals with high versus low levels of signature 2 mutations (Figure 3 C,D). Second, we obtained methylation data for a testis cell line, produced by the Encyclopedia of DNA Elements (ENCODE) project [22]. Signature 2 mutations are ~8.5 times as likely to occur in regions of high (>=50%) versus low (<50%) methylation. We do not detect any difference in this ratio between regions, or between individuals with high versus low signature 2 rates, although the number of mutations involved is probably too low to provide much power (Methods; Fisher’s exact test P=0.14). Third, we tested dependence on *B* statistic [23], a measure of conservation. We found that the relative magnitudes of both signatures 1 and 2 depend on *B* statistic, but that both these dependencies were independent of the per-population intensities of the signatures (Figure 4 A,B). This, along with a similar result for recombination rate, (Figure 4 C,D) confirm that these differences are not strongly associated with differences between population in patterns of selection, recombination, or recombination-related processes such as gene conversion.

**Figure 3:**
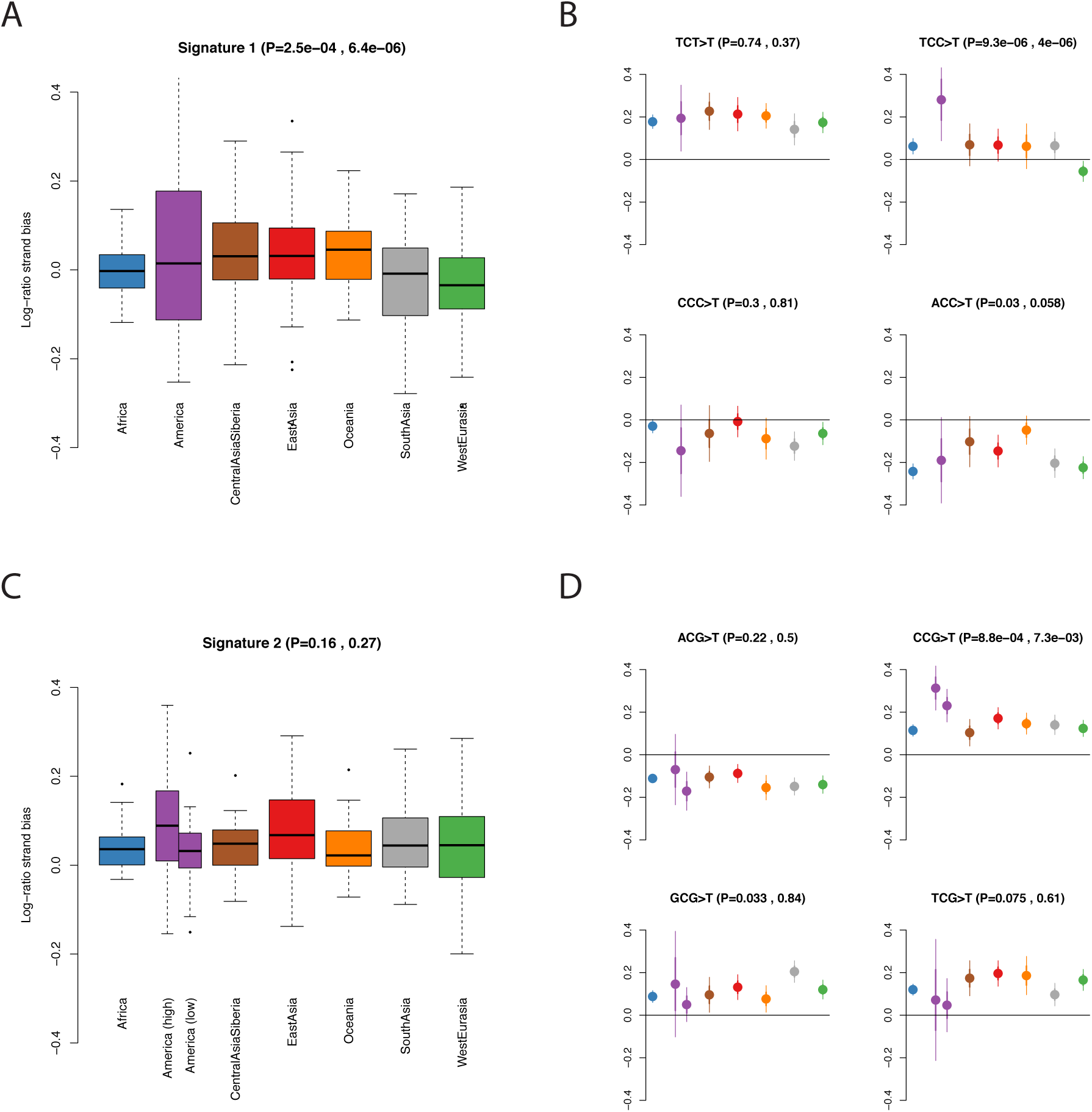
Transcriptional strand bias in mutational signatures. We plot the log of the ratio of *f*_2_ mutations occurring on the untranscribed versus transcribed strand. Therefore a positive value indicates that the C>T mutation is more common than the G>A mutation on the untranscribed (i.e. coding) strand. P values in brackets are, respectively, ANOVA P-values for a difference between regions and t-test P-values for a difference between **i)** West Eurasia and other regions (excluding South Asia) in A&B **ii)** 11American samples with high rates of signature 2 mutations and other regions in C&D. **A**: Boxplot of per-individual strand bias for mutations in signature 1 (TCT>T, TCC>T, CCC>T and ACC>T). One sample (S_Mayan-2) with an extreme value (0.48) is not shown. **B**: Population-level means for each of the mutations comprising signature 1. **C,D**: as A&B but for signature 2. We separated out the 11 American samples with high rates of signature 2 mutations.

**Figure 4:**
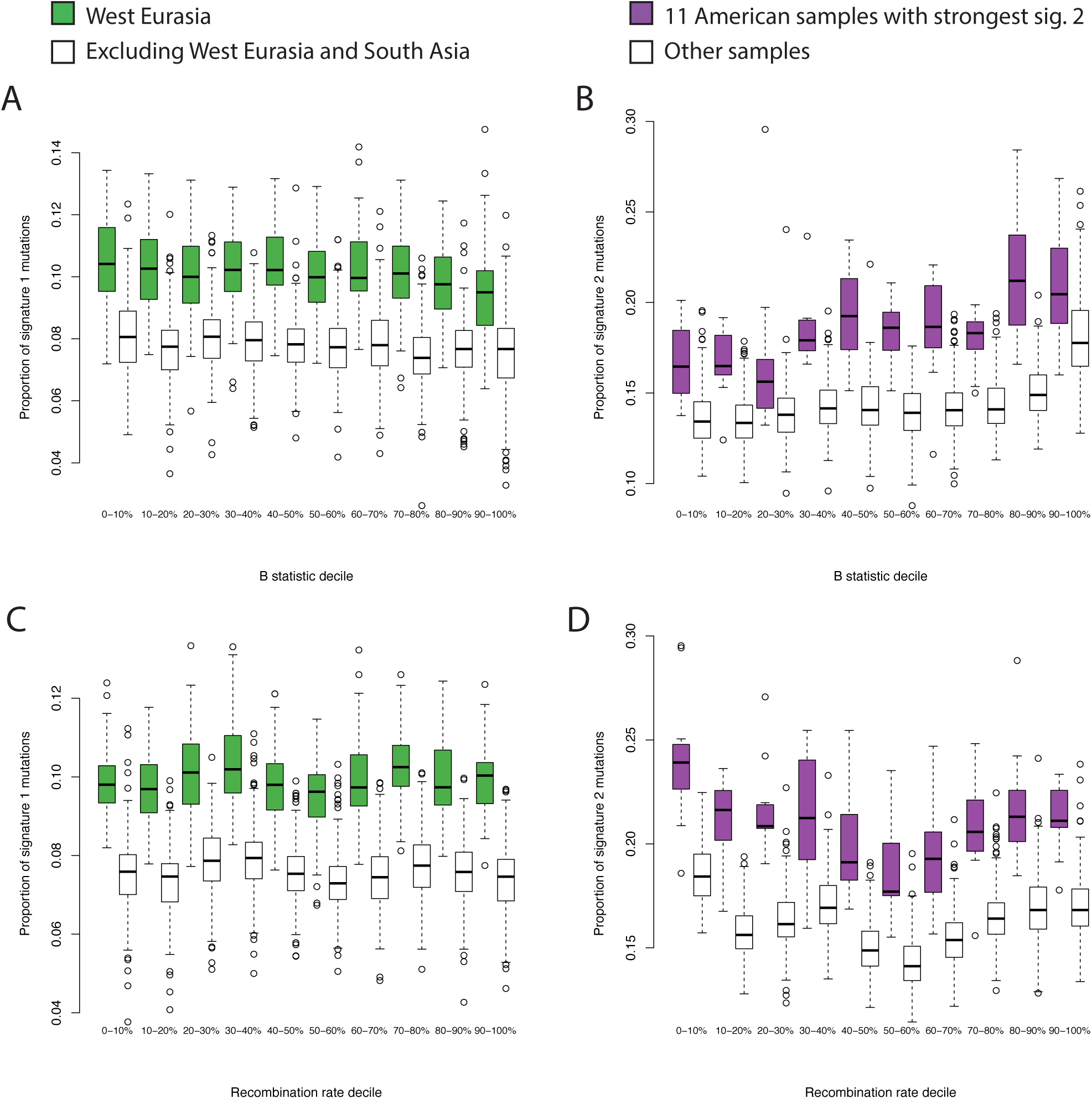
Dependence of signatures on genomic features. **A,B**: dependence on conservation, measured by *B* statistic (0=lowest *B*statistic; highest conservation). **A**: Comparison of proportions of signature 1 mutations between West Eurasia and other populations (excluding South Asia). **B**: Comparison of proportions of signature 2 mutations between the 11 American samples with the highest proportions, and all other samples. **C,D**: As A&B, but showing dependence on recombination rate decile computed in 1kb bins.

Most of the variation in signature 2, however, can be explained by differences in demography between populations (Figure 5). In particular, a relatively high proportion of signature 2 mutations are repeat mutations (i.e. mutations that have occurred more than once in different individuals), and the frequency spectrum of such mutations is more sensitive to demography–particularly recent expansions–than non-repeat mutations. To show this, we first looked at the proportion of variants at different frequencies that were in signature 2 (i.e. C->T mutations at CpG sites; Figure 5A). There is a strong enrichment of these variants in Native Americans at frequency 2, but not for singletons, nor for frequencies greater than 3. It is hard to imagine a purely mutational process that would affect variants of frequency 2, but not 1. Next, instead of restricting to variants of a particular frequency, we counted the proportion of derived alleles per genome that are in signature 2 (Figure 5B, Methods). While there is an increase in this proportion in Native Americans, it is extremely small – an increase in proportion of 1.6×10^−5^ relative to East Asians. Further, this increase is not restricted to Native Americans with high rates of signature 2 *f*_2_ mutations. This suggests that while there may be subtle variation in the rate of signature 2 mutations, the effects we observed are not driven by this, but rather by the fact that signature 2 mutations have been shifted into different frequency classes in different populations, relative to other mutations. One important property of signature 2 mutations is that CpG sites have a much higher mutation rate than nonCpG sites [24], and therefore a much higher rate of repeat mutations. For example, ~12% of *de novo* CpG mutations are expected to occur at sites that are already polymorphic in 1000 Genomes phase 1 (n=1,092) [25], and 87% of exonic *de novo* CpG mutations are polymorphic in ExAC (n=60,706) [26] – rates that are about ten times higher than those for non-CpG mutations. In the SGDP (n=300), 17.7% of signature 2 *f*_2_ muations are shared between Africans and non-Africans, compared to 8.3% of all *f*_2_ mutations, suggesting that around 9% of signature 2 *f*_2_ mutations are repeat mutations. This shifts the relative frequency spectra because the spectrum of repeat mutations is more sensitive to recent population growth than that of non-repeat mutations (a similar argument applies for triallelic sites [27]). In particular, under recent population growth genealogies become more star-like and the numbers of singleton non-repeat mutations increases, but the number of doubleton repeat mutations increases even more (Figure 5C). This means that the ratio of CpG to non-CpG variants at any given frequency is extremely sensitive to recent demography, and the patterns that we observe could be explained by recent exponential growth on the order of between 10- and 100- fold in most populations (Figure 5D). Thus, it seems likely that differences in the proportion of rare, or private, variants in this class is driven by differences in the rate of recent population growth rather than differences in mutation rate and implies that Native American populations with high rates of rare signature 2 mutations experienced rapid population growth after the initial founding bottleneck of the Americas.

**Figure 5:**
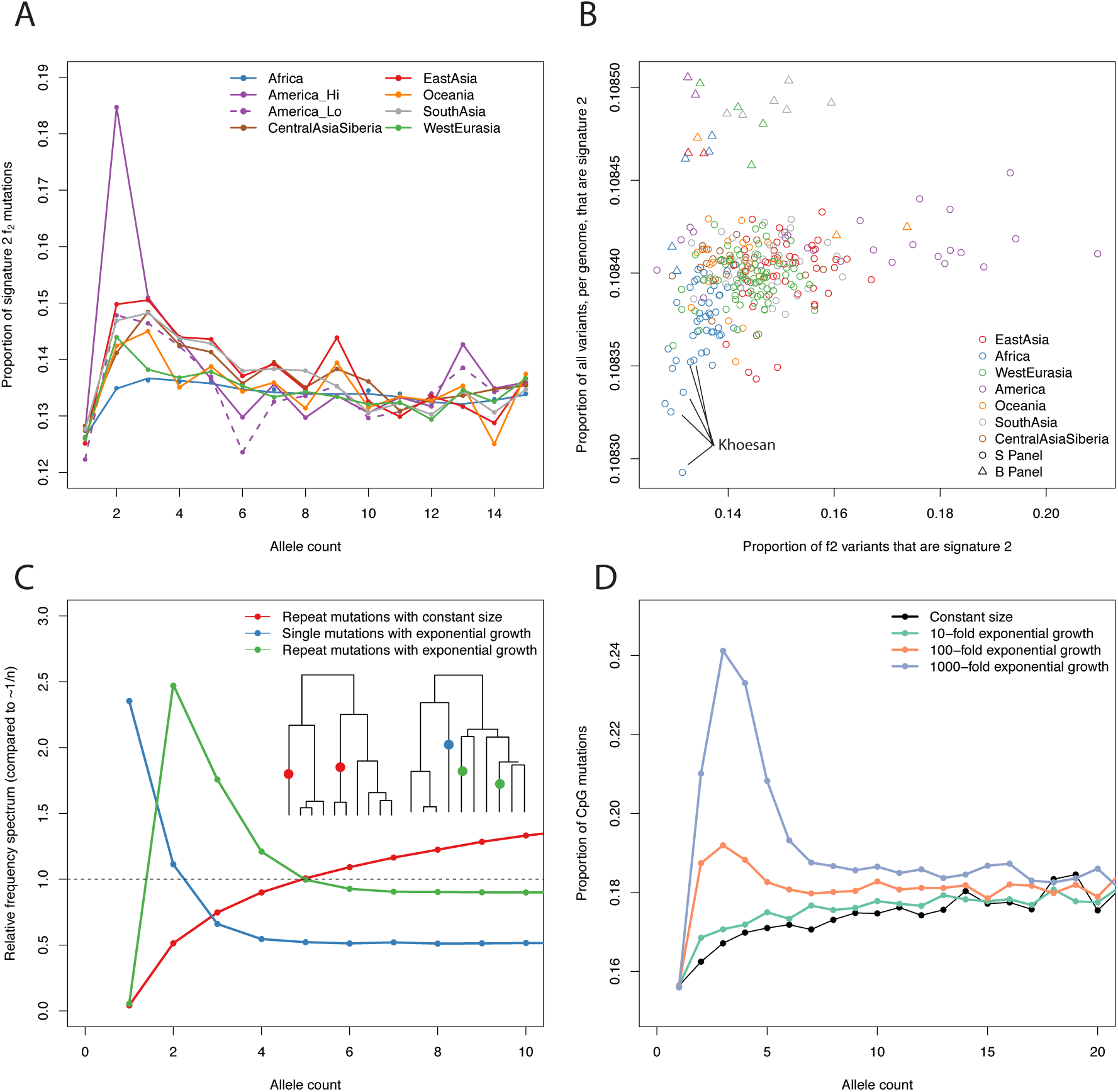
Differences in signature 2 can be explained by demography. **A**: The proportion of variants that are in signature 2 for different regions, for allele counts from 1 to 20. **B**: The proportion of variants that are in signature 2 for *f*_2_ variants on the x-axis, and all variants per-genome on the y-axis. Samples in SGDP panel B, processed in a different pipeline, shown as triangles. **C**: Simulated allele frequency spectra for repeat mutations for 50 haplotypes under the standard (i.e. constant population size) coalescent, and both single and repeat mutations under the coalescent with exponential growth (100-fold in 0.04 *Ne*generations). The y-axis is scaled by the expected frequency of single mutations in the constant size case (i.e. 1/n). Inset trees show examples of the genealogies obtained – constant size on left, exponential growth on right. Results from 200,000 independent trees. **D**: Simulation of the proportion of mutations that are at CpG sites at different frequencies, assuming that 15% of all mutations are CpGs and 10% of CpGs are repeat mutations. Compare to **A**.

In contrast, differences in signature 1 are consistent with a difference in mutation rate. In particular, individuals with a high rate of signature 1 *f*_2_ variants also have a high total proportion of signature 1 mutations (Figure 6A), and we see enrichment in Europeans relative to other groups in singletons, and for variants with allele counts up to around 30, corresponding to a frequency of around 5% (Figure 6B). The enrichment changes as a function of frequency, which suggests that the increase in mutation rate might have changed over time. Therefore, to study the time depth of these signals, we investigated whether signature 1 could be detected in ancient samples by constructing a corrected statistic, that measures the intensity of the mutations enriched in signature 1, normalized to reduce spurious signals that arise from ancient DNA damage (methods). This statistic is enriched to present-day European levels in both an eight thousand year old European hunter-gatherer and a seven thousand year old Early European Farmer [28] but not in a 45,000 year old Siberian [29], nor in the Neanderthal [30] or Denisovan [31] genomes (Figure 6C) – consistent with a recent estimate that this increase in mutation rate lasted between 2,000 and 15,000 years before present [9]. The statistic is predicted by neither estimated hunter-gatherer ancestry, nor early farmer ancestry, in 31 samples from 13 populations for which ancestry estimates were available [28] (linear regression p-values 0.22 and 0.15, respectively). Thus the effect is not strongly driven by this division of ancestry. If it has an environmental basis, it is not predicted by latitude (linear regression of signature 1 loadings against latitude for West Eurasian samples; p=0.68), but is predicted by longitude (p=6×10^−8^; increasing east to west).

**Figure 6:**
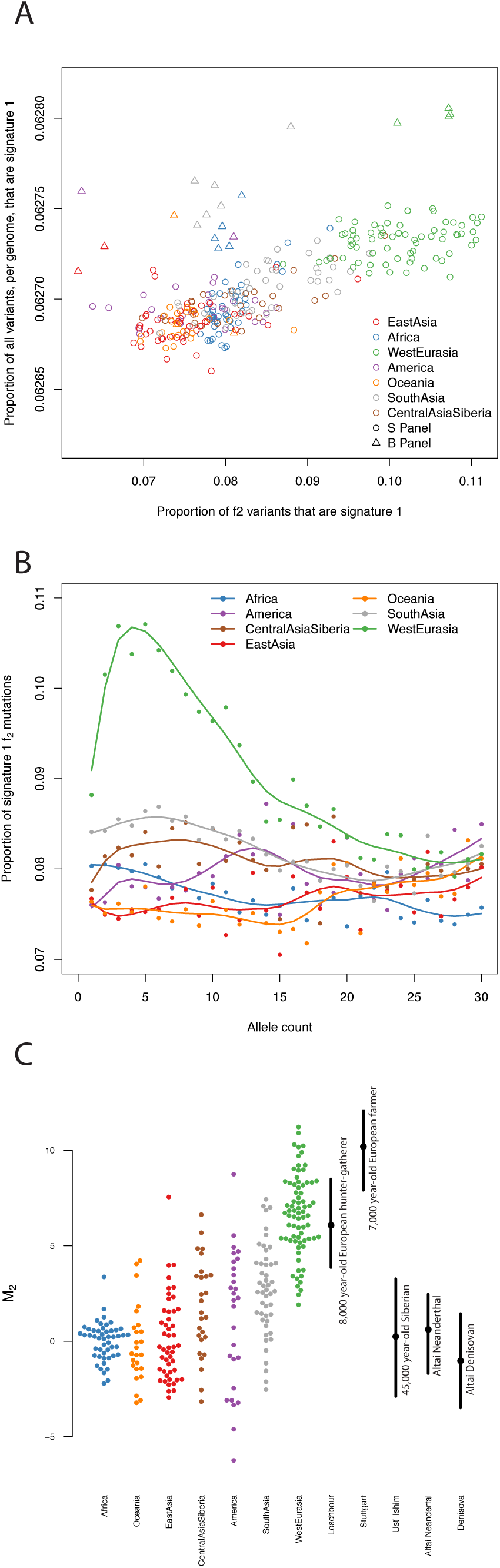
Details of signature 1 **A**: The proportion of variants that are in signature 1 for *f*_2_ variants on the x-axis, and all variants per421 genome on the y-axis. Samples in panel B, processed in a different pipeline, shown as triangles. **B**: Proportion of mutations in signature 1 as a function of derived allele count from 2 to 30. **C**: Signature 1, corrected to be robust to ancient DNA damage (Methods), for *f*_2_ variants in the SGDP and five high coverage ancient genomes. Solid lines show 5-95% bootstrap quantiles.

## Discussion

We characterized two independent differences among human populations in their spectrum of rare variants, however this may not be comprehensive. Our power to detect differences in variation spectra depends on a number of factors, including sample size, and the level of background variation. While modest differences in variant spectra might be much more widespread than we describe here [9], it is clear that the West Eurasian signature 1 enrichment is by far the most dramatic. Two questions naturally follow from this result. First, does this result imply a difference in absolute mutation rate? And second, what is the biological basis behind this signature?

In our previous analysis of the SGDP data [13] we showed that the rate of mutation accumulation differed between populations. In particular, mutation accumulation, relative to chimpanzee, was consistently around 0.1% higher in non-Khoesan groups than Khoesan groups, and around 0.5% higher in non-Africans than Africans. Since the mean divergence time between two humans is much less than the mean divergence between humans and chimp, these results imply a much greater difference in mutation rate – for example we estimated that the rate of mutation accumulation would be around 5% higher on the non-African relative to the non-African branch. The proportion of f2 mutations attributable to signature 1 (i.e TCT>T, TCC>T, CCC>T and ACC>T) increases from a mean of 7.8% in Africans to 10.0% (range 8.8-11.1%) in West Eurasians. If we make the assumptions that the only differences in mutation rate are the ones we detected, the absolute rates of all other mutation types are the same between populations, and the difference in mutation rate has been present for the entire period since the divergence of Africans and non-Africans, then this change implies a maximum increase in genome-wide mutation rate of 2.3% (range 1.1-3.6%). This is in insufficient to explain the approximately 5% excess of mutations in West Eurasian in the SGDP data, and is also likely to be a large overestimate of the possible effect since Harris and Pritchard suggest that the elevated rate of mutation accumulation in this class was largely restricted to 15,000 to 2,000 years ago instead of persisting over the whole period since the divergence of Africans and non-Africans [9]. In any case, as we previously observed [13], this cannot explain the difference in total mutation accumulation rate, because that effect is not restricted to West Eurasians.

We cannot be definitive about the biological cause of variation in signature 1, but our analyses provide a clue. In terms of the immediate mutagenic cause, signature 1 is most similar to COSMIC [19] signature 11 (Pearson correlation *ρ*=0.81), which is associated with alkylating agents used as chemotherapy drugs, damaging DNA through guanine methylation. The reversal of transcriptional strand bias for this signature in West Eurasians supports the idea that the increased rate of these mutations in West Eurasians is driven by damage to guanine bases, consistent with deamination of methyl-guanine to adenine, leading to the G>A (equivalently C>T) mutations that we observe. An increase in this rate might be driven by an increase in guanine methylation, either through environmental exposure, or through inherited variation that affected demethylation pathways. Signature 1 is also highly correlated with COSMIC signature 7 (*ρ*=0.75), caused by ultraviolet (UV) radiation exposure but it is difficult to imagine how this could affect the germline, would not explain our observed increase in ACC>T mutations, would not be expected to reverse the strand bias, and should produce an enrichment of CC>TT dinucleotide mutations in West Eurasians that we do not observe (p=0.41). Harris (2015) [8] suggested that UV might cause germline mutations indirectly through folate deficiency in populations with light skin pigmentation (since folate can be degraded in skin by UV radiation). It is unknown what mutational signature would be caused by this effect, but the fact that we do not observe enrichment of signature 1 in other lightly pigmented populations like Siberians and northeast Asians suggests that it is not driving the signal.

Our analysis of signature 2 underscores the importance of modeling repeat mutations, at least for CpG sites, in rare variant analysis. One consequence is that any analysis that restricts to part of the frequency spectrum is potentially confounded by this effect – this includes subtle effects that might arise from studies that have differential power to call rare variants among samples – implying that it might be difficult to reliably detect differences in CpG mutation rate from polymorphism data. Nonetheless it seems that the relative rate of CpG mutation accumulation does vary across populations, but only very slightly. Our results also suggest that the CpG:non-CpG ratio as a function of frequency could be a useful statistic for estimating the rate of recent population growth and that some Native American populations have experienced extremely rapid growth in recent history.

It is important to understand changes in the mutation rate on the timescale of hominin evolution in order to calibrate demographic models of human evolution [32] and the observation of variation in mutation spectra *between* populations [8] made this calibration even more complicated. Further work in this area will involve more detailed measurement of mutation rates in diverse populations – to date, most work on somatic, cancer, or *de novo* germline mutations has been conducted in populations of West Eurasian origin – and the extension of these approaches to other populations will be required to fully understand variation in mutation rates and its consequences for demographic modeling.

## Methods

### Identifying mutational signatures

We used SNPs called in 300 individuals from the Simons Genome Diversity Project [13] (SGDP). The SGDP provides position- and sample-specific masks, with strictness ranging from 0-9 (0=least strict). We first called variants at filter level 1, independently in each individual, which is recommended for most analyses. This gave us a list of sites that were reliably variable in at least one of the 300 samples. Then, to avoid underestimating the frequency of variants due to some samples being masked, we recalled all these sites in every individual at the less strict filter level 0. We polarized SNPs assuming that the chimpanzee reference panTro2 carried the ancestral allele (ignoring sites where the chimp genome could not be aligned to the human genome), and classified by the two flanking bases in the human reference (hg19). We restricted to sites of given derived allele counts. For example, when we analyze *f*_2_ variants, we consider both variants where a single individual is homozygous derived and variants where any two separate individuals (ignoring population labels) are heterozygous derived. We count two mutations if the individual is homozygous and one if it is heterozygous. We then merged reverse complement classes to give counts of SNPs occurring in 96 possible mutational classes. Finally, we normalized these counts by the frequency of ATA>ACA mutations. The remaining matrix represents the normalized intensity of each mutation class in each sample, relative to the sample with the lowest intensity. Formally, let *C*_*ij*_ be the counts of mutations in class *i* for sample *j*. Then, the intensities that we analyze, *X*_*ij*_ are given by,

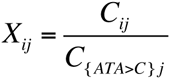

We decomposed this matrix ***X*** using non-negative matrix factorization [15] implemented in the *NMF R* package [16] with the multiplicative algorithm introduced by Lee & Seung [15], initialized using the non-negative components from the output of a *fastICA* analysis [33] implemented in the *fastICA* package in R (https://cran.r-project.org/web/packages/fastICA/index.html). For the diagnostic plots in Supplementary Fig. 2, we used 200 random starting points to compare the results of different runs. When we initialized the matrix randomly, rather than using *fastICA*, we obtained a slightly closer fit to the data (root-mean-squared error in ***X*** of 0.024 vs 0.025) and similar factor distributions (Supplementary Fig. 6A), except that all signatures were dominated by CpG mutations (Supplementary Fig. 6B). Removing a constant amount of each CpG mutation from each signature recovered signatures closer to the *fastICA*-initialized signatures (Supplementary Fig. 6C), so we concluded that this was a model-fitting artifact, and did not reflect true signatures. Finally we performed the analysis on a matrix normalized be the total number of mutations in each sample 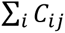 rather than the number of ATA>C mutations. (Supplementary Figure 6).

The ordering of the factors is arbitrary so, where necessary, we reordered for interpretability. To plot mutational signatures and compare with the COSMIC signatures, we rescaled the intensities of each class according to the trinucleotide frequencies in the human reference genome. The scale of the weightings is therefore not easily interpretable. To perform principal component analysis on ***X***, we normalized so that the variance of each row was equal to 1.

### Analysis of 1000 Genomes data

We classified 1000 Genomes variants according to the ancestral allele inferred by the 1000 Genomes project, and counted the number of *f*_2_ and *f*_3_ variants carried by each individual in each mutation class. We ignored SNPs that were multi-allelic or where the ancestral state was not confidently assigned (confident assignment denoted by a capital letter in the “AA” tag in the “INFO” field of the vcf file). For each individual, we computed the proportion of the total mutations carried by that individual that were in each of signatures 1 and 2. We excluded the five outlying samples: HG01149 (CLM), NA20582 & NA20540 (TSI), NA12275 (CEU), NA19728 (MXL) which had extreme values in one of these signatures.

### Transcriptional strand

We downloaded the knownGenes table of the UCSC genes track from the UCSC genome browser (http://genome.ucsc.edu/). Taking the union of all transcripts in this table, we classified each base of the genome according to whether it was transcribed on the + or – strand, both, or neither (including uncalled bases). These regions totaled 607Mb, 637Mb, 36Mb and 1,599Mb of sequence respectively. We then counted mutations (not collapsed with their reverse complements) in our dataset that occurred in regions that were transcribed on the + or – strand, ignoring regions where both or neither strand was transcribed.

### Methylation status

We downloaded the Testis_BC 1 and 2 (two technical replicates from the same sample) tables from the HAIB Methyl RRBS track from the UCSC genome browser (http://genome.ucsc.edu/). We constructed a list of 33,305 sites where both replicates had >=50% methylation and another list of 166,873 sites where both replicates had <50% methylation. We then classified the CpG mutations in our dataset according to which, if either, of these lists they fell into. Ultimately, there were only 1186 classified mutations in the whole dataset, including 43 in Native American samples and 12 in Native American samples with high rates of signature 2. Therefore, although we found no significant interactions between methylation status and population, it may be simply that we lack power to detect it.

### *B* statistic and recombination rate

We classified each base of the genome according to which decile of *B* statistic [23] or HapMap 2 combined recombination rate [34] (in 1kb blocks) it fell into and counted mutations in each class.

### Analysis of total number of mutations

To count the total mutations per-genome in Figures 5A and 6A, we counted mutations at all frequencies, rather than restricting to variants at a particular frequency in the whole dataset. We excluded the last 20Mb of chromosome 2, where 46 samples had high rates of missing data.

### Coalescent simulations

We simulated a sample of 50 haplotypes under the standard coalescent, by first simulating a coalescent tree, and then generating mutations on the tree as a Poisson process. For the simulations shown in Figure 5, we simulated 200,000 independent trees. To simulate repeat mutations, we simulated two mutations and performed an OR operation on the genotype vectors – this correctly captures the probabilities of nested and non-nested mutations. To simulate exponential growth, we first simulate under the standard coalescent, and then rescale time *t*such that the new time *t’* is given by:

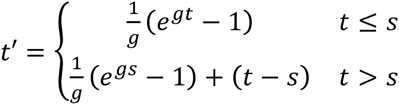

where 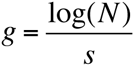 to simulate *N*-fold growth starting at time s. We simulated for *N*=100 and 1000 and chose *s*=0.01 in coalescent time, corresponding to 0.01×2N_e_ generations, or around 9,000 years if we assume human-like parameters of N_e_=15,000 and a generation time of 30 years.

### Analysis of ancient genomes

We identified heterozygous sites in five ancient genomes from published vcf files, and restricted to sites where there was a single heterozygote in the SGDP. The corrected signature 1 log-ratio is defined by

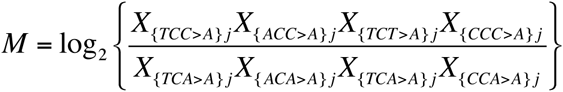

and then normalized so that the distribution in African populations has mean 0 and standard deviation 1. We estimated bootstrap quantiles by resampling the counts *C*_*ij*_ for the ancient samples and recomputing *M*.

### Increase in absolute mutation rate

Suppose that in a single sample there are *M* mutations in total, of which *N* are from a particular signature. Let 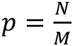. Suppose the number of mutations in that signature increases by Δ*N*, but the number of all other mutations stays the same. Then the new proportion of mutations in the signature is 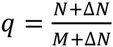. Under these assumptions, the increase in the total mutation rate 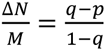.

## Code availability

Scripts used to run the analysis are available from https://github.com/mathii/spectrum.

## Acknowledgments

We thank Mark Lipson, Priya Moorjani and Swapan Mallick for helpful comments. I.M. is supported by a long-term fellowship from the Human Frontier Science Program LT001095/2014-L. D.R. is supported by NIH grant GM100233 and is a Howard Hughes Medical Institute Investigator.

---

**Supplementary Figure 1:**
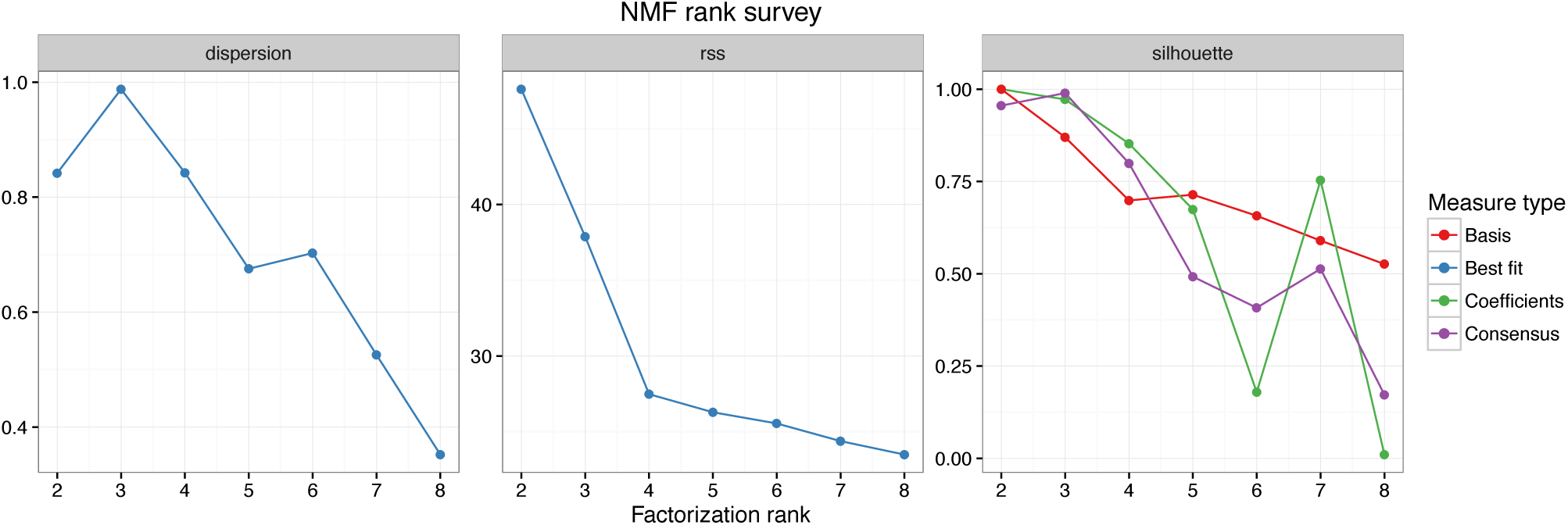
Diagnostic plots for NMF using variants of frequency 2. Each plot shows the value of a measure, computed over 50 random start points, for factorization ranks from 2 to 8. From left to right: Dispersion, a measure of reproducibility of clusters across runs (1=perfectly reproducible); Residual sum of squares (lower=better fit); Silhouette, a measure of how reliably elements can be assigned to clusters (1=perfectly reliably).

**Supplementary Figure 2:**
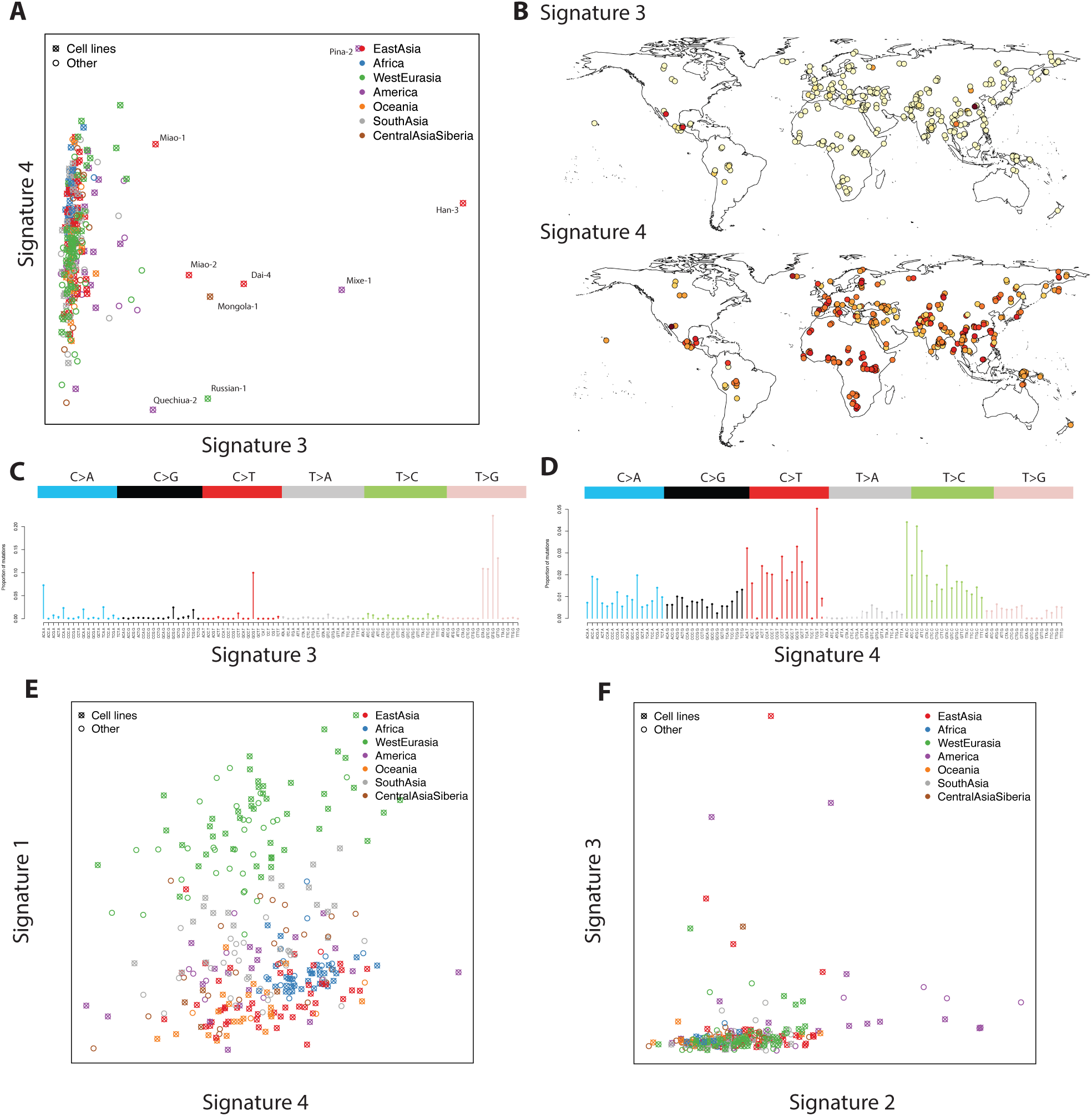
Distribution and characterization of mutational signatures2,4 3 and 4. **A**: Per-sample coefficients for signatures 3 and 4. **B**: Geographic distribution of signatures 3 and 4. **C**: Mutational spectrum of signature 3. **D**: Mutational spectrum of signature 4. **E-F**: Comparison of loadings of 1 and 2 with signatures 3 and 4. In supplementary plots, we denote the signatures obtained from *f_r_*variants with rank *k* by signature_r,k_, so that signature_2,4_ is equivalent to the signature in the main text.

**Supplementary Figure 3:**
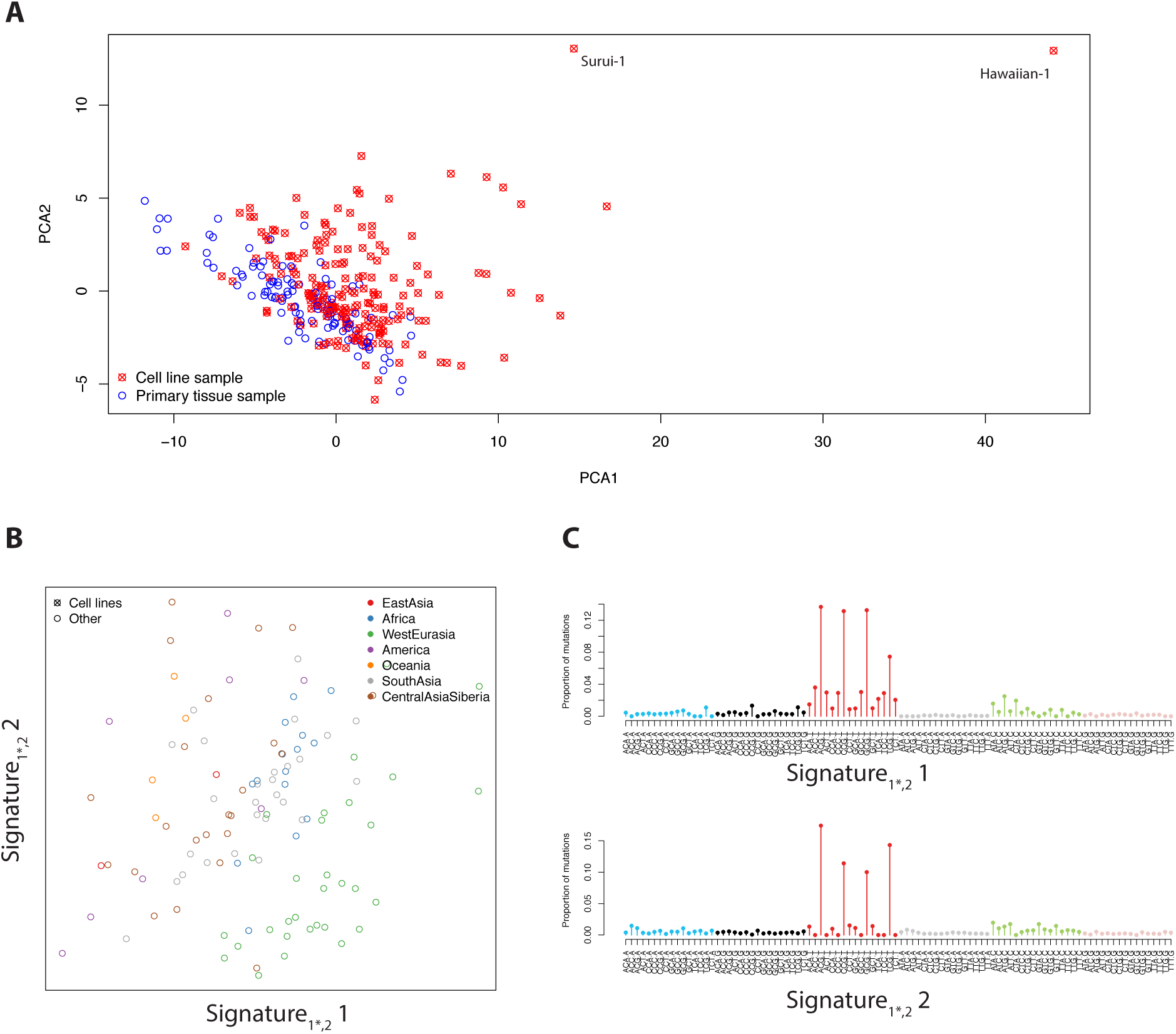
Analysis of *f*_1_ variants **A**: The first two principal components of the mutational spectrum of *f*_1_ variants, showing the difference between cell line and primary tissue derived samples. **B**&**C**: Mutational signatures inferred from *f*_1_ variants with rank 2, but excluding cell line samples. **B**: Factor loadings for signature_1*,2_ 1 and 2 (asterisk denotes no cell lines). **C**: Mutational signatures_1*,2_ 1 and 2. Signature_1*,2_ 1 is confounded with CpG mutations in this case, but clearly shows an elevated level of TCC>T and ACC>T mutations.

**Supplementary Figure 4:**
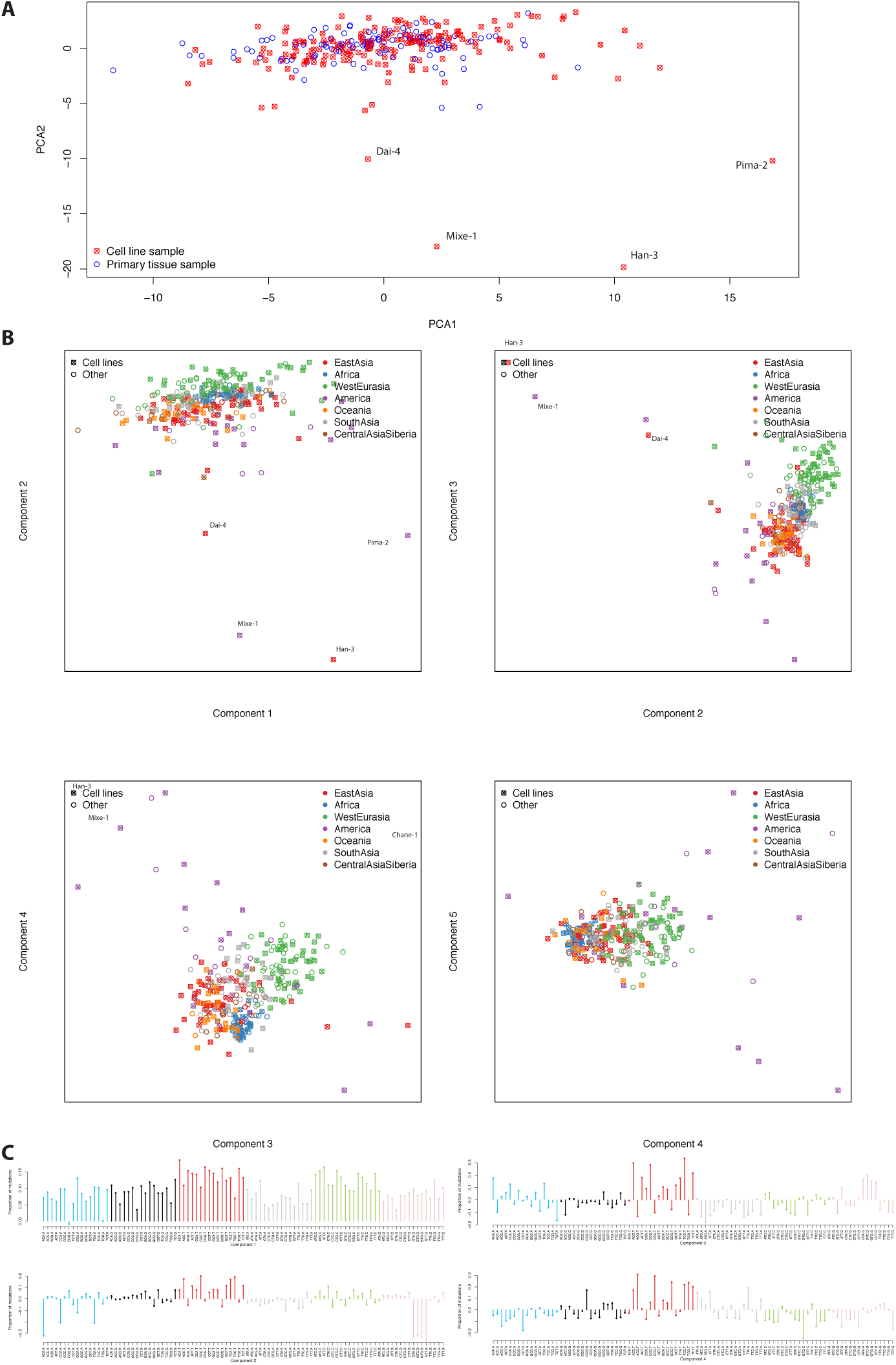
Principal component analysis of the mutational spectrum of f_2_ variants. **A:** The first two principal components of the mutational spectrum of *f*_2_ variants, showing no difference between cell line and primary tissue derived samples. **B**: Principal component positions. Labeled by sample source (A) and geographic region (B). **C**: Component loadings. Note that principal components2,3 and 4 correspond roughly to mutational signatures_2,4_ 3, 2 and 1 respectively.

**Supplementary Figure 5:**
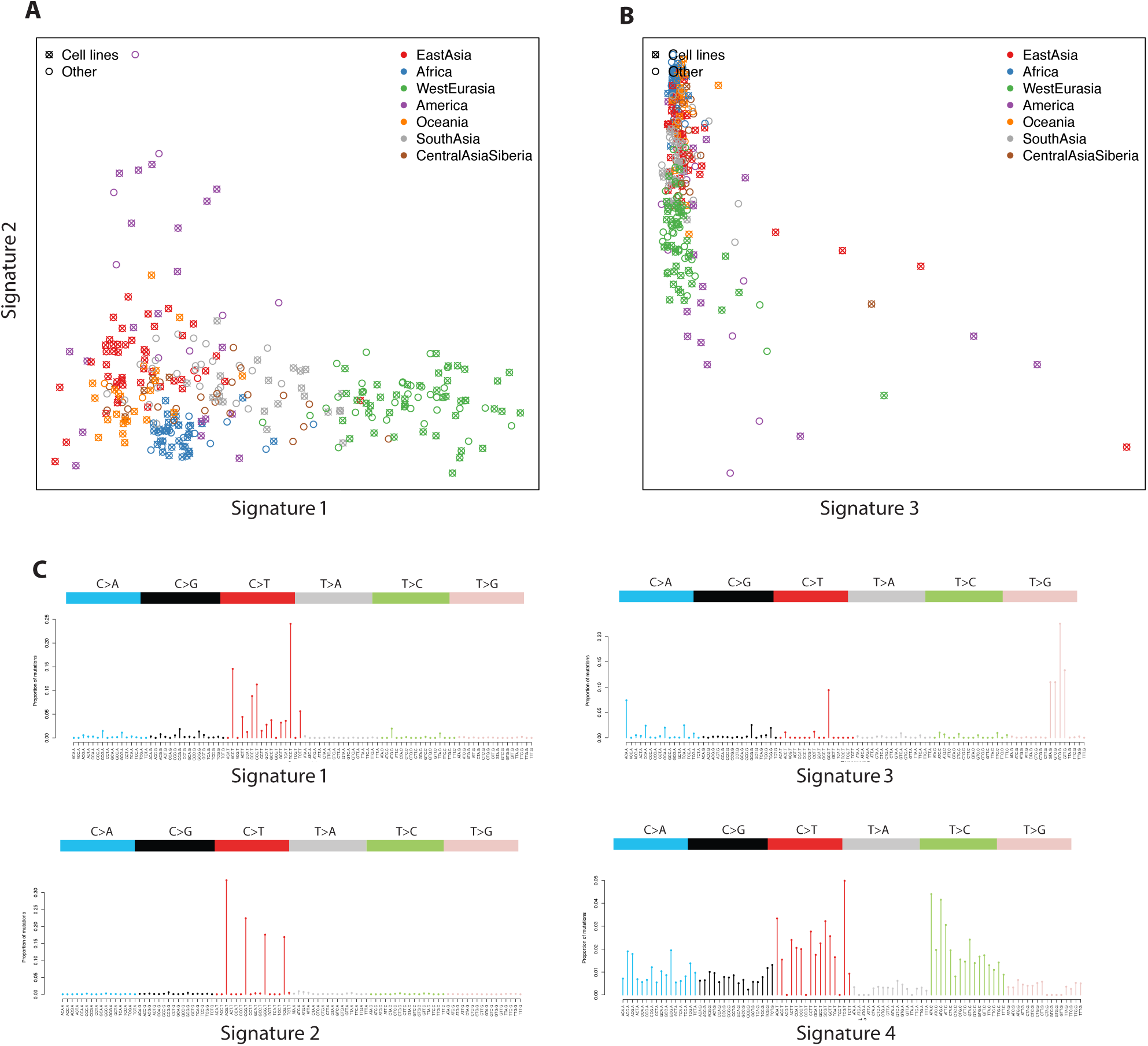
NMF analysis of f2 variants at rank 4 - as the main analysis, 555 but normalizing the mutational spectra by the total number of mutations in each sample, rather than the number of ATA>C mutations.

**Supplementary Figure 6:**
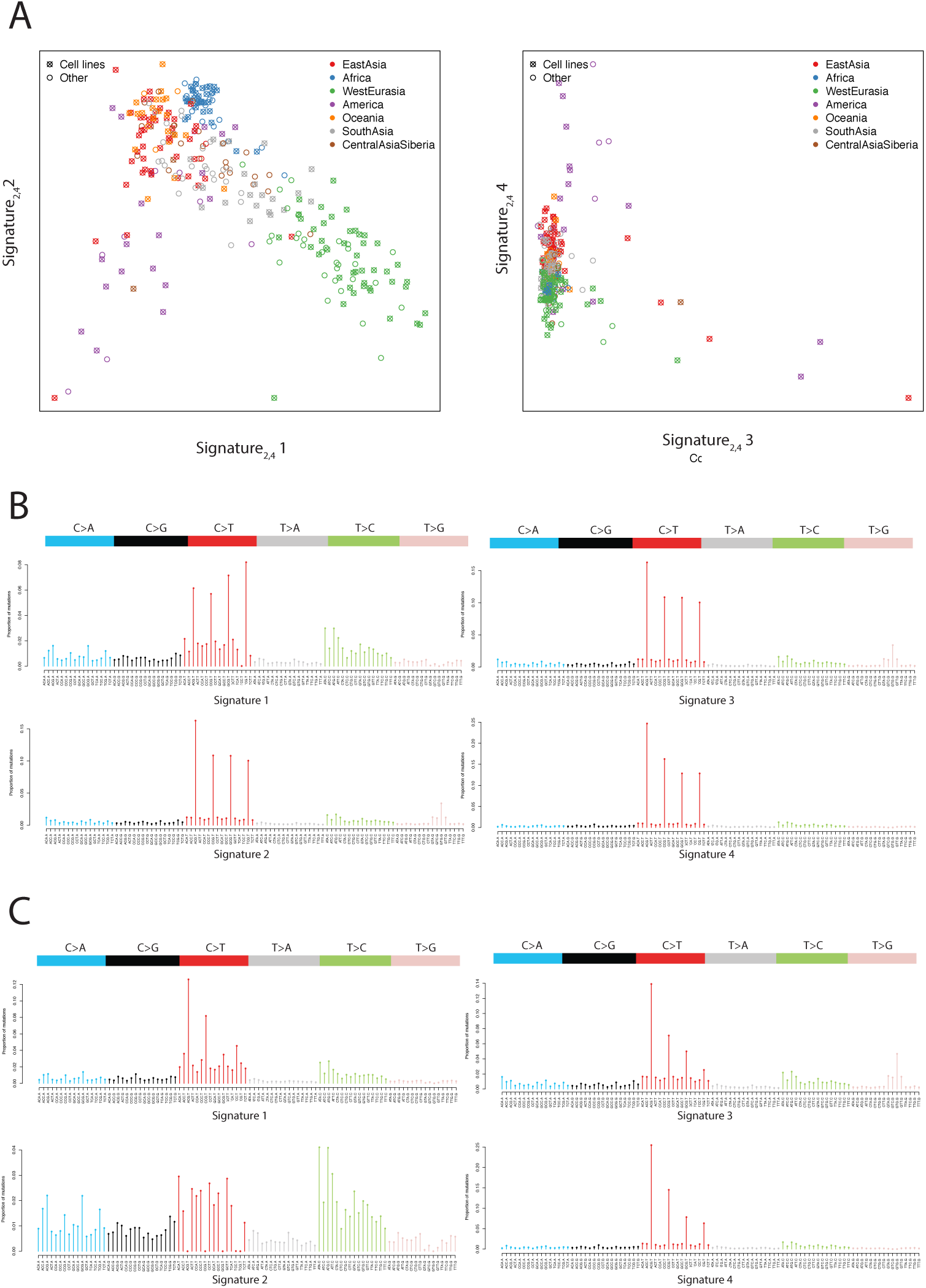
NMF analysis of *f*_2_ variants at rank 4 with random initialization of the NMF algorithm. **A**: Distribution of signatures across samples. **B**: Mutational signatures 1-4. **C**: Mutational signatures 1-4 where, for each CpG mutation class, we subtracted the minimum over all four signatures from the signature.

